# Cassava HapMap: Masking deleterious mutations in a clonal crop species

**DOI:** 10.1101/077123

**Authors:** Punna Ramu, Williams Esuma, Robert Kawuki, Ismail Y Rabbi, Chiedozie Egesi, Jessen V Bredeson, Rebecca S Bart, Janu Verma, Edward S Buckler, Fei Lu

## Abstract

Cassava (*Manihot esculenta* Crantz) is an important staple food crop in Africa and South America, however, ubiquitous deleterious mutations may severely reduce its fitness. To evaluate these deleterious mutations in the cassava genome, we constructed a cassava haplotype map using deep sequencing from 241 diverse accessions and identified over 28 million segregating variants. We found that, 1) while domestication modified starch and ketone metabolism pathways for human consumption, the concomitant bottleneck and clonal propagation resulted in a large proportion of fixed deleterious amino acid changes, raised the number of deleterious mutations by 26%, and shifted the mutational burden towards common variants; 2) deleterious mutations are ineffectively purged due to limited recombination in cassava genome; 3) recent breeding efforts maintained the yield by masking the most damaging recessive mutations in the heterozygous state, but unable to purge the mutation burden, which should be a key target for future cassava breeding.

---

Cassava is the third most consumed carbohydrate source for millions of people in tropics, after rice and maize^1^. Even though cassava was domesticated in Latin America^2^, it has spread widely and become a major staple crop in Africa. Cassava stores starch in underground storage roots, which remain fresh until harvest. Cassava is a highly heterozygous species. Although its wild progenitor, *M. esculenta* ssp. *falbellifolia*, reproduces by seed^3^, it is particularly worth noted that cultivated cassava is almost exclusively clonally propagated via stem cutting, in which a single individual contributes its entire genome to its offspring^4^. The limited number of recombination events in such vegetatively propagated crops results in a potential accumulation of deleterious mutations across the genome^5^. Thus, mutation burden in cassava is expected to be more severe than in sexually propagated species. Deleterious mutations are considered to be at the heart of inbreeding depression^6^. Inbreeding depression is extremely severe, even in elite cassava accessions, where a single generation of inbreeding results in >60% reduction in fresh root yield^7,8^. In this study, we aimed to identify deleterious mutations in cassava populations, which in turn can help accelerate cassava breeding by allowing breeders to purge deleterious mutations more efficiently.

We conducted a comprehensive characterization of genetic variation by whole genome sequencing (WGS) of 241 cassava accessions, including 203 elite breeding accessions (*M. esculenta* Crantz), 16 close relatives (*M. esculenta*. ssp. *flabellifolia, M. esculenta* ssp. *peruviana*) of modern cultivars^2,9^, 11 hybrid/tree cassava accessions, and 11 more divergent wild relatives (*M. glaziovii* and others) (**Supplementary Fig. 1** and **Supplementary Table 1**). Samples included 54 accessions from an initial haplotype map I (HapMapI) study^10^. Wild *M. glaziovii* has been used extensively in cassava breeding programs to transfer disease resistance alleles to cultivated cassava (e.g., Amani Breeding program)^8^. On average, more than 30x coverage sequences were generated for each accession. The 518.5 Mb cassava genome (v6.1) has roughly 51% repetitive elements with several common recent retrotransposons^10^. To exclude misalignment and ensure high quality of variant calling, repeat sequences were pre-filtered using repeat bait (**Supplementary Fig. 2**) and the remaining sequences were aligned against the cassava reference genome v6.1^10,11^. Variants from low copy regions of the genome were identified to develop the cassava haplotype map II (HapMapII) with 27.8 million variants (25.9 million SNPs and 1.9 million indels) and with a low error rate of 0.01%, which is the proportion of segregating sites in the reference accession (**Supplementary Fig. 3**). The correlation between read depth and proportion of heterozygotes of SNPs is extremely low (*r^2^* = 6E-05, **Supplementary Fig. 4**). Cultivated cassava exhibited 9.94 million variants (**Supplementary Table 2**), of which nearly 50% were found to be rare (<5% minor allele frequency (MAF)) (**Supplementary Table 2 and Supplementary Fig. 5**). Haplotypes were phased and missing genotypes were imputed with high accuracy using BEAGLE v4.1^12^ (accuracy *r*^2^ = 0.966) (**Supplementary Fig. 6**). Linkage disequilibrium was as low as in maize^13^ and decayed to an average *r^2^* = 0.1 in 3,000 bp (**Supplementary Fig. 7**).

Cultivated cassava presented lower nucleotide diversity (π = 0.0036) compared with its progenitors (*M. esc. ssp. flabellifolia*, π = 0.0051). In addition, a close relationship between the two species was observed from phylogenetic analysis (**Supplementary Fig. 8**). Both lines of evidence support the hypothesis that cultivated cassava was domesticated from *M. esc. ssp. flabellifolia*^2,9,10^. To evaluate population differentiation of cassava, a principal component (PC) analysis was performed and showed substantial differentiation among all cassava species and hybrids **(Fig. 1a)**, where cultivated cassava showed moderate genetic differentiation from its progenitors (*F_st_*: 0.16), and high genetic differentiation from tree cassava (*F_st_*:0.32) and wild relatives (*F_st_*:0.44) (**Supplementary Table 2 and Supplementary Figs. 9 and 10**). However, PC analysis showed very little differentiation among cultivated cassava **(Fig. 1b)**, where geographic subpopulations of cultivated cassava presented surprisingly low value of *F_st_* among themselves (0.01-0.05) despite the fact that these subpopulations were sampled from different continents (**Supplementary Table 2**). This suggests that despite clonal propagation, there has been enough crossing to keep cultivated cassava in one breeding pool.

**Figure 1.**
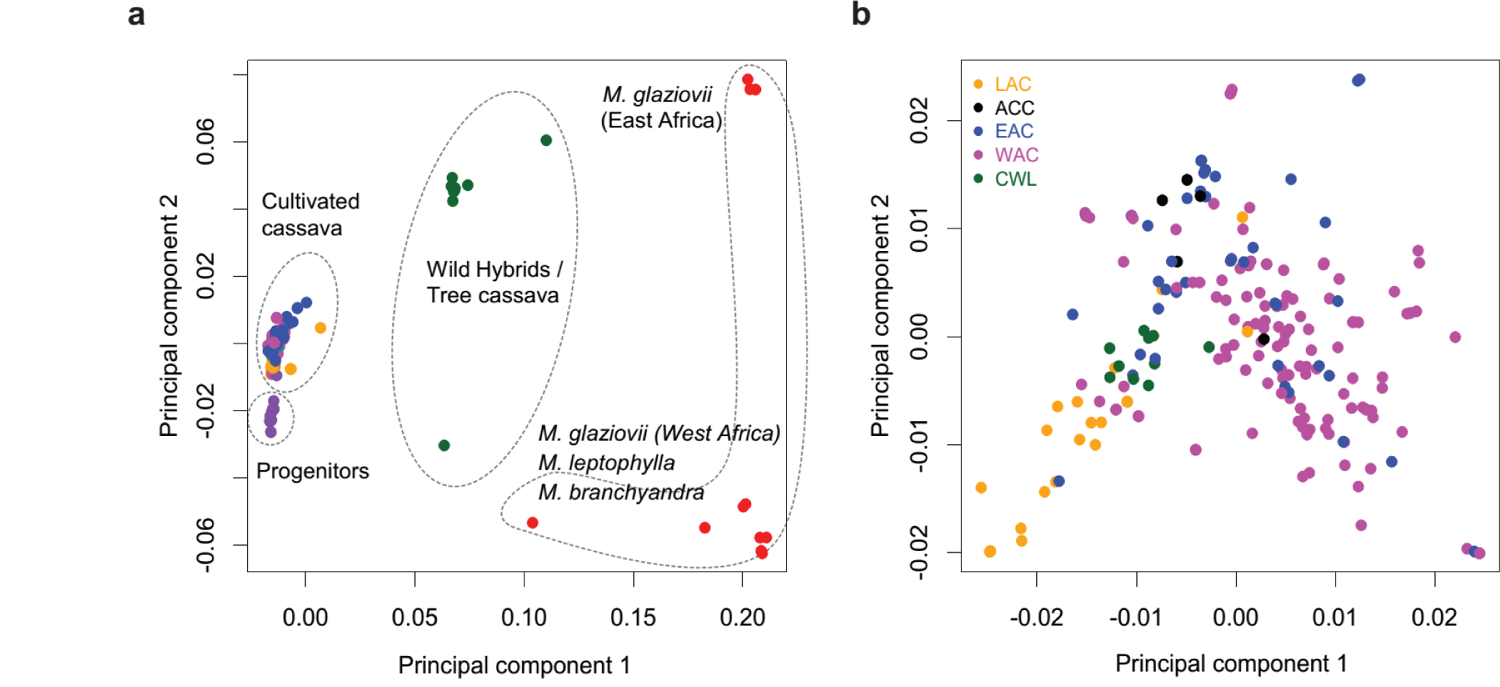
Principal component analysis (PCA) of cassava accessions included in cassava HapMapII. (a) PCA of all cassava accessions (progenitors, cultivated, and wild cassava accessions). A total of 43.8% genetic variance is captured in first two principal components. (b) PCA of cultivated cassava clones. A total of 9.1% genetic variance is captured in first two principal components. The abbreviations are represented as follows: LAC – Latin American cassava, ACC – Asian Cultivated cassava, EAC – East African cassava, WAC – West African cassava, CWL – Crosses between WAC and LAC.

Sequence conservation is a powerful tool to discover functional variation^14,15^. We identified deleterious mutations by utilizing genomic evolution and amino acid conservation modeling. The cassava genome was aligned to seven species in the Malpighiales clade to identify evolutionarily constrained regions of cassava genome. Based on genomic evolutionary rate profiling (GERP)^16^ score, nearly 104-Mb of the genome (20%) of cassava was constrained (GERP score > 0) (**Supplementary Fig. 11**). The evolutionarily constrained genome of cassava (104 Mb) is comparable to maize (111 Mb)^17^ in size, but less than humans (214 Mb)^16^ and more than *Drosophila* (88 Mb)^18^. GERP profiling also identified remarkably asymmetric distribution of constrained sequence at the chromosome scale (**Supplementary Fig. 12**). In addition to the constraint estimation at the DNA level, consequences of mutation on amino acids in proteins were assessed using Sorting Intolerant From Tolerant (SIFT) program^19^. Nearly 3.0% of coding SNPs in cultivated cassava were non-synonymous mutations (**Supplementary Table 2** and **Supplementary Fig. 13**), of which 19.3% (57,952) were putatively deleterious (SIFT < 0.05). As the strength of functional prediction methods varies^14^, we combined SIFT (< 0.05) and GERP (> 2) to obtain a more conservative set of 22,495 deleterious mutations (**Supplementary Fig. 14**).

To estimate the individual mutation burden, we used rubber (*Hevea brasiliensis*), which diverged from the cassava lineage 27 million years ago^10^, as an out-group to identify derived deleterious alleles in cassava. First, we focused on the fixed deleterious mutations. The derived allele frequency (DAF) spectrum shows that cassava (5%, **Fig. (2)** appears to have more fixed deleterious mutations than maize (3.2%, DAF > 0.8)^20^ when compared at the same threshold (SIFT < 0.05). Across cultivated cassava there were 150 fixed deleterious mutations. These deleterious mutations cannot be purged through standard breeding which relies on recombination of segregating alleles, but these fixed deleterious mutations are the potential targets for genome editing^21^. Together with the other 22,345 segregating deleterious mutations, the mutation burden in cassava was substantial. Given the several millennia of breeding in the species, why are these deleterious mutations still in cultivated cassava and how were breeders managing them? We evaluated the effects of recombination, selection, and drift, as the main processes controlling the distribution of deleterious mutations in the genome.

**Figure 2.**
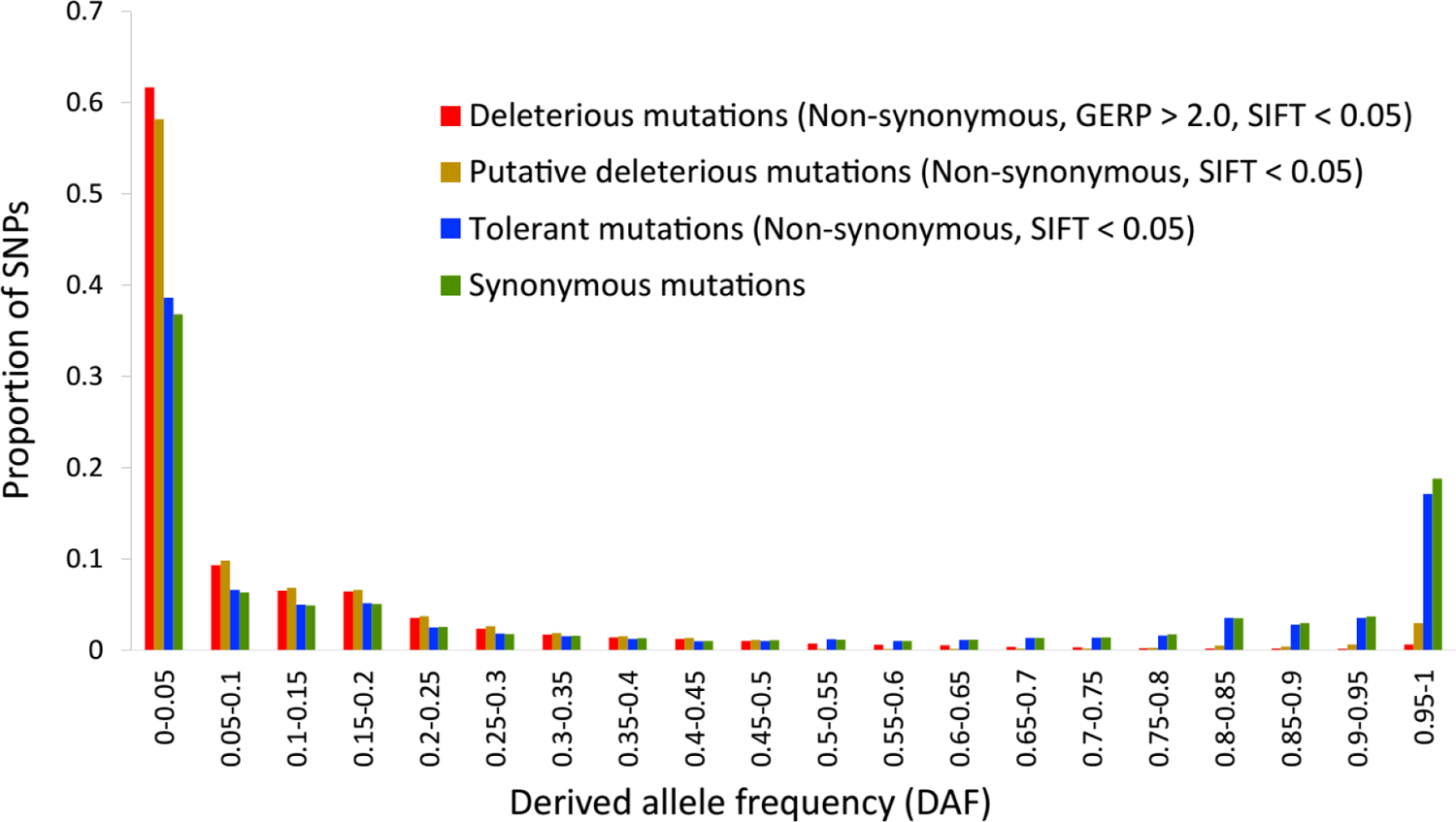
Site allele frequency spectrum of deleterious mutations in cassava genome. Derived allele frequency (DAF) distribution of alleles are presented. Rubber genome is used as the out group to define derived alleles.

Recombination is an essential process to purge deleterious mutations from genome^22^. In vegetatively propagated species like cassava, recombination is expected be less efficient in purging deleterious mutations. This hypothesis was supported by a weak correlation between recombination rate and distribution of deleterious mutations (*r* = -0.065, *P* = 0.13, **Fig. 3a**). Deleterious mutation were nearly uniformly spread across the cassava genome (**Fig. 3b and Supplementary Fig. 15**), rather than being concentrated in low recombination regions as in human^23^, fruit fly^24^, and maize^17^. Thus, recombination, which is presumably rare in a clonally propagated crop, does not effectively purge mutation burden in cassava.

**Figure 3.**
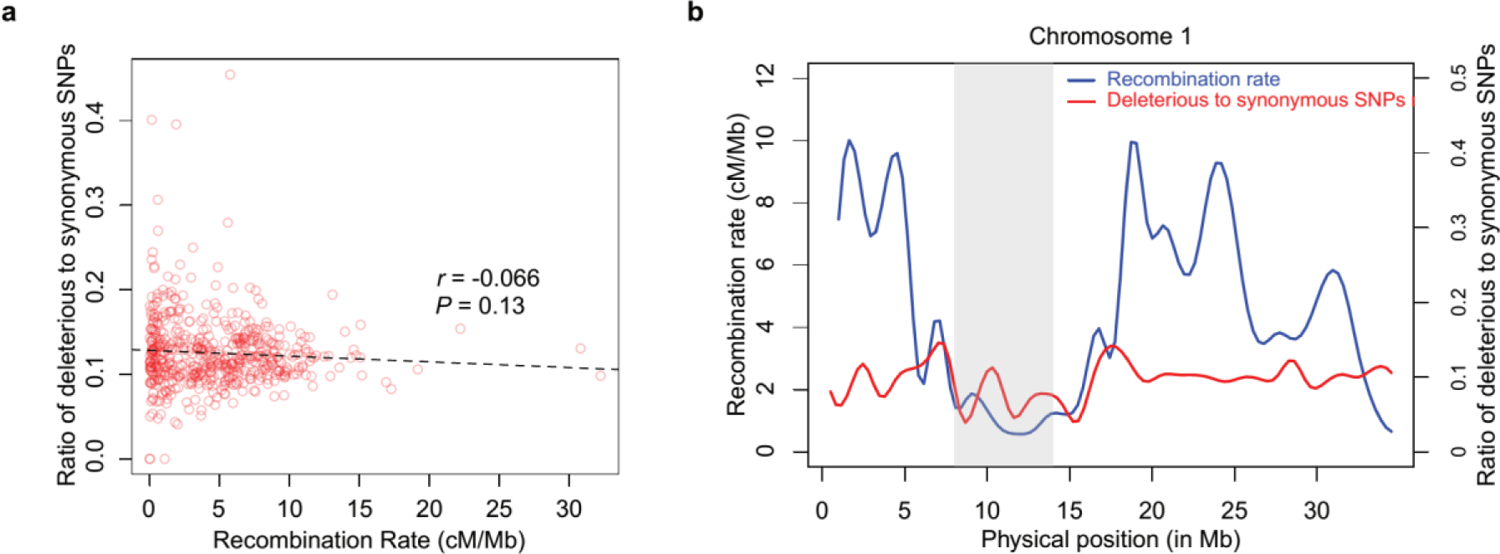
Effect of recombination on the distribution of deleterious mutations in cassava genome. (a) Correlation between recombination rate and number of deleterious mutations in the genome. (b) Distribution of deleterious mutations as a function of recombination rate on chromosome 1.

Domestication is important in evolution and improvement of crop species. The major domestication trait of cassava is the large carbohydrate rich storage root. Cultivated cassava has 5-6 times higher starch content than its progenitor^3^. Another domestication trait is the reduced cyanide content in roots^3^. Every tissue of cassava contains cyanogenic glucosides^25^. Ketones, cyanohydrin, and hydrogen cyanide are the key toxic compounds formed upon degradation of cyanogenic glucosides^25,26^. These toxic compounds have to be eliminated before consumption. To identify the genomic regions under selection during the domestication, a likelihood method (the cross-population composite likelihood ratio, XP-CLR)^27^ was used to scan the genome in Latin American accessions and the progenitor *M. esculenta*. ssp. *flabellifolia*. We identified 203 selective sweeps containing 427 genes in Latin American accessions (**Supplementary Fig. 16a**). Genes in these sweep regions were enriched for starch and sucrose synthesis (3.8-fold enrichment; FDR = 7.2 × 10^-03^) and cellular ketone metabolism (3.4-fold enrichment; FDR = 5.3 × 10^-03^) (**Supplementary Fig. 16b**). The results suggest that selection during domestication increased production of carbohydrates and reduced cyanogenic glucoside in cassava. Likewise, selection signatures of recent bottleneck event in African cassava accessions were also evaluated. A total of 244 selective sweeps were identified containing 416 genes. These genes were enriched for serine family amino acid metabolism (4.2-fold enrichment, FDR = 2.1 × 10^-06^) and cellular response to stress (1.3-fold enrichment, FDR = 4.9 × 10^-06^, **Supplementary Fig. 17**). Since L-Serine is involved in the plant response to biotic and abiotic stresses^28,29^, together with the functional enrichment of cellular response to stress, it may reflect that disease resistance accessions were selected in recent breeding program in Africa^8^.

How was the genetic burden shaped in the selective sweeps? We found that Latin American accessions showed 25% less (*P* = 0.009, **Fig. 4a**) deleterious mutations than progenitors in sweep regions. Similarly, African accessions exhibited a 35% drop (*P* = 2.1 × 10^-07^, **Fig. 4b**) in sweeps compared to Latin American accessions. In addition to the comparison between populations, significant reductions of deleterious mutations were observed within population by comparing sweep regions and the rest of the genome. For example, selective sweeps presented 44% depletion (*P* = 9.7 × 10^-12^, **Fig. 4c**) of deleterious mutations in Latin American accessions and 41% reduction (*P* = 8.7 × 10^-130^, **Fig. 4d**) in African accessions. This implies that haplotypes containing fewer deleterious alleles were favored during selection.

**Figure 4.**
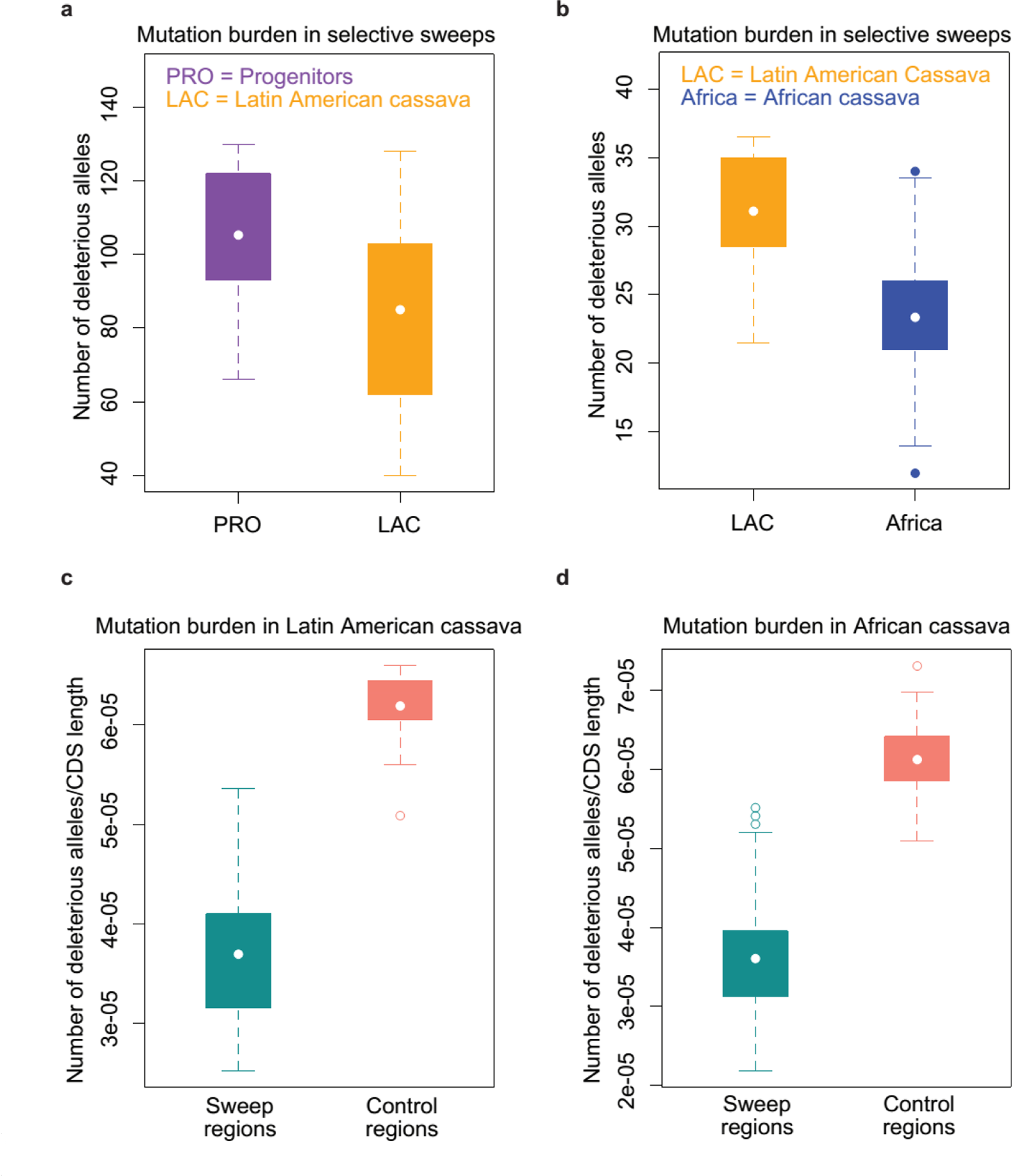
Mutation burden in selective sweep regions. (a) Mutation burden between progenitors and Latin American cassava accessions in domestication sweep regions. (b) Mutation burden between Africa and Latin American cassava accessions in sweep regions identified in recent improvement in Africa. (c) Mutation burden in Latin American cassava accessions between domestication selective sweeps and control regions (rest of the genome). (d) Mutation burden in African cassava accessions between sweep regions identified in recent improvement and control regions (rest of the genome) in Africa.

However, drift after domestication played a more important role in affecting mutation burden in cassava. Although Latin American accessions and African accessions had a similar number of deleterious mutations (*P* = 0.42, **Fig. 5a**), they presented a prominent increase of total burden by 26% (*P* = 9.1 × 10^-09^, **Fig. 5a**) when compared with progenitors, and shifted the mutation burden towards common deleterious variants (**Supplementary Fig. 18**). The increase of deleterious mutations during domestication was also found in dog^30^. The results suggest that the severe bottleneck of domestication and shift from sexual reproduction to clonal propagation resulted in a rapid accumulation of deleterious mutations in cultivated cassava.

**Figure 5.**
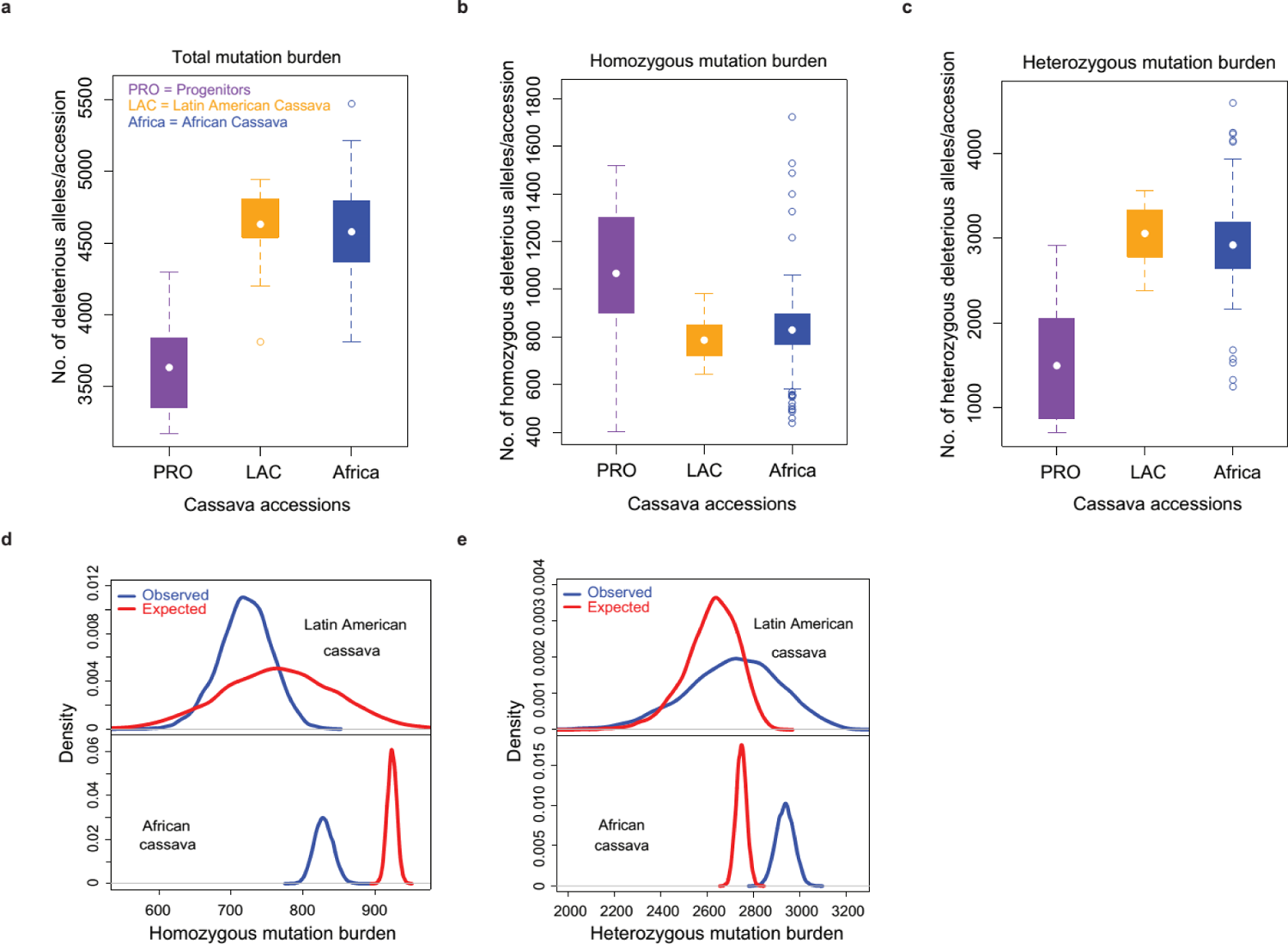
Mutation burden in cassava populations. (a) Total mutation burden in progenitors, Latin American cassava and African cassava accessions. Bottleneck during domestication increased mutation burden. Demography in Africa has no significant influence on mutation burden in African cassava accessions. (b) Homozygous mutation burden in cassava populations. Domestication decreased homozygous mutation burden in cultivated cassava. (c) Heterozygous mutation burden in cassava populations. Domestication increased heterozygous mutation burden in cultivated cassava. (d) Comparison between the observed homozygous mutation burden and the expected homozygous mutation burden under HWE assumption in cultivated cassava. (e) Comparison between the observed heterozygous mutation burden and the expected heterozygous mutation burden under HWE assumption in cultivated cassava.

How have the breeders been able to maintain yield, given the substantial growth of mutation burden in cultivated cassava? This became apparent when the homozygous deleterious mutations and heterozygous deleterious mutations were compared. Relative to *M. esculenta*. ssp. *flabellifolia*, the homozygous mutation burden substantially decreased by 23% (*P* = 7 × 10^-03^, **Fig. 5b**) in cultivated accessions regardless of the elevated frequency of deleterious alleles (**Supplementary Fig. 18**), while the heterozygous mutation burden remarkably increased by 96% (*P* = 8.1 × 10^-07^, **Fig. 5c**), despite the reduced genetic diversity in cultivated cassava (π = 0.0036) relative to progenitors (π = 0.0051). In addition to the comparison between cultivated cassava and progenitors, we also compared observed and expected mutation burden under the assumption of Hardy-Weinberg Equilibrium (HWE) within cultivated cassava (**Online Methods**). Although HWE was probably never reached in the breeding pool, the relative depletion of homozygous mutation burden and excess of heterozygous mutation burden would not be seen unless it was selected and maintained. The results from bootstrap resampling (10,000 times) showed that the observed homozygous mutation burden was less than the expected (Latin American cassava: 5.6% decrease, *P* = 0; African cassava: 10.3% decrease, *P* = 0, **Fig. 5d**), and the observed heterozygous mutation burden was more than expected (Latin American cassava: 3.5% increase, *P* = 1.5 × 10^-312^; African cassava: 6.9% increase, *P* = 0, **Fig. 5e**), indicating a significant deviation from HWE expectation. These evidence suggest that breeders have been trying to manage the recessive deleterious mutations in the heterozygous state to mask their harmful effects.

Mutations with large homozygous effect are more likely to be recessive^31^. We found nearly 64.5% of deleterious mutations occurred only in the heterozygous state (**Supplementary Fig. 19**). Although the low allele frequency confines effective tests for the excess heterozygotes of these deleterious mutations, they are more likely to be strong deleterious mutations, resulting in the significant yield loss in the first generation of selfed cassava plants^7,8^. These mutations were in genes (n = 7,774) mainly enriched for macromolecule catabolism and biosynthesis (**Supplementary Fig. 20a**). In contrast, the deleterious mutations existing predominantly in the homozygous state (proportion of homozygotes > 70%, **Supplementary Fig. 19**), were present in genes (n = 245) enriched for amine and ketone metabolism, as well as chemical and stimulus responses (**Supplementary Fig. 20b**). This difference suggests that the deleterious mutations primarily exhibited in the heterozygous state may have relatively large fitness consequences.

Cassava is a major staple crop feeding hundreds of millions of people. Using deep sequencing of a comprehensive and representative collection of 241 cassava accessions, we developed the HapMapII, a highly valuable resource for cassava genetic studies and breeding. In this vegetatively propagated species, deleterious mutations have been accumulating rapidly due to the lack of recombination. The bottleneck event during domestication exacerbated the existing mutation burden in cassava. Breeding efforts successfully maintained the yield by selecting high fitness haplotypes at a few hundred loci and handling most damaging mutations in the heterozygous state. However, breeders were unable to purge the mutation burden due to limited recombination, instead they shielded deleterious mutations by increasing the heterozygosity while screening thousands of potential hybrids (**Supplementary Fig. 21**). In the short term, this practice for managing mutation burden may produce gains in yield. In the long run, however, a mutational meltdown may be triggered by new mutations, decreasing genetic diversity in breeding pool, and clonal propagation. The deleterious mutations should be important targets for future cassava breeding programs. Genomic selection and genomic editing technologies^21^ are anticipated to help purge deleterious mutations and improve this globally important crop.

## ONLINE METHODS

### Samples and whole genome sequencing

To maximize the diversity and representation for cassava, all samples were selected based on breeders’ choice and diversity analysis from accessions included in Next Generation Cassava Breeding project (www.nextgencassava.org). Whole genome sequences were generated from 241 cassava accessions including 203 elite breeding accessions, 16 progenitors (*M. falbellifolia, M. peruviana*)^7^, 11 hybrid/tree cassava accessions and 11 wild relative cassava accessions (*M. glaziovii* and others) (**Supplementary Table 1**). Among 241 cassava accessions, 172 accessions were sequenced at the Genomic Diversity Facility at Cornell University, Ithaca, NY, USA. Standard Illumina PCR-free libraries were constructed with insert size of 500-bp using Illumina standard protocol. Sequences of 200-bp length were generated using Illumina HiSeq 2500 and 150-bp length were generated using NextSeq Series Desktop sequencers. Donald Danforth Plant Science Center, St. Louis, MO, USA generated ~20x coverage sequences for 15 elite cassava accessions. Sequences for remaining 54 cassava accessions were collected from HapMapI^10^, generated at the University of California at Berkeley (USA).

### Alignment of reads and variant calling for generation of cassava haplotype map (HapMapII)

The cassava genome was found to have large amounts of repeat sequences^10^. To minimize misalignment, these repeats were pre-filtered by aligning the sequences to a bait containing repeat sequences and organelle sequences (**Supplementary Fig. 1**). Remaining sequences after pre-filtering were aligned to reference genome (v6.1) using burrows-wheeler alignment with maximal exact matches (BWA-MEM) algorithm (http://bio-bwa.sourceforge.net/bwa.shtml#13). To ensure high quality SNP calling, especially for those rare variants, we developed an in-house pipeline, FastCall (https://github.com/Fei-Lu/FastCall), to perform the stringent variant discovery. The procedures include: 1) Genomic positions having both insertion and deletion variants were ignored, since these sites were likely in complex regions with many misalignments; 2) For multiple allelic sites, if the third allele had more than 20% depth in any individual, the site was ignored; 3) For a specific site, if the minor allele did not have a depth between 40% and 60% in at least one individual when individual depth was greater than 5, the site was ignored; 4) A chi square test for allele segregation^13^ in all individual is performed. The sites with *P*-value more than 1.0 × 10^-03^ were ignored. 5) On average, over 30X depth was used to for individual genotype calls. The genotype likelihood was calculated based on multinomial test reported by Hohenlohe *et. al*^32^. To remove potential spurious variants arising from paralogs, an additional filter was applied to keep only variants with depth between 7,500 and 11,500. The missing data was about 4%. The genotypes were imputed and phased into haplotypes using BEAGLE v4.1^12^. A total of 10% of the genotypes were masked before imputation to calculate the imputation accuracy.

### Population genetics analysis

SNP density, pair-wise nucleotide diversity (π), Tajima’s D and *F_st_* were calculated using VCFtools^33^ (**Supplementary Fig. 8**). Principal component analysis was carried out in Trait Analysis by aSSociation, Evolution and Linkage (TASSEL)^34^. Recombination rates were obtained from cassava HapMapI source^10^.

### Genomic evolutionary rate profiling (GERP)

Constrained portion of cassava genome was identified by quantifying rejected substitutions (strength of purifying selection) using GERP++ program^16^. Multiple whole genome sequence alignment was carried out for the seven species in Malpighiales clade of plant kingdom, including cassava, rubber (*Hevea brasiliensis*), jatropha (*Jatropha curcas*), castor bean (*Ricinus communis*), willow (*Salix purpurea*), flax (*Linum usitatissimum)*, and poplar (*Populus trichocarpa)*. Phylogenetic tree and neutral branch length (estimated from 4-fold degenerate sites) were used to quantify constraint intensity at every position on cassava genome. Cassava genome sequence was eliminated during the site specific observed estimates (RS scores) to eliminate the confounding influence of deleterious derived alleles segregating in cassava populations that are present in reference sequence.

### Identifying deleterious mutation

Amino acid substitution and their effects on protein function were predicted using ‘Sorting Tolerant From Intolerant (SIFT)’ algorithm^19^. Non-synonymous mutations with SIFT score < 0.05 were defined as putative deleterious mutations. SIFT (< 0.05) and GERP (>2) annotations were combined to identify the deleterious mutations existing in constrained portion of the genome. These deleterious mutations were used to calculate mutation burden of cassava.

### Identifying selective sweep regions

Cross-population composite likelihood approach (XP-CLR) method^27^ was used to identify the selective sweeps in two contrasts: Latin America cassava accessions (test populations) against progenitors (*M. esc*. ssp *flabellifolia*, reference population) for domestication event and African cassava accessions (test populations) against Latin American cassava accessions (reference population) for recent improvement in Africa. Selection scan was performed across the genome using 0.5 cM sliding window between the SNPs spacing of 2-kb. A genetic map of cassava generated by International Cassava Genetic Map Consortium^35^ was used in the XP-CLR analysis. XP-CLR scores were normalized using Z-score and smoothed spline technique with R-package (GenWin)^36^. Outlier peaks were selected which were above than 99 percentile of normalized values. AgriGO^37^ and REVIGO^38^ tools were used for GO enrichment analysis.

### Mutation burden in cassava accessions

Number of derived deleterious alleles present in each cassava accessions were counted to identify mutation burden in cassava accessions in three models (homozygous mutation burden, heterozygous mutation burden, and total mutation burden). Homozygous mutation burden is the number of derived deleterious alleles in the homozygous state. Heterozygous mutation burden is the number of derived deleterious alleles existing in the heterozygous state. Total mutation burden is the number of derived deleterious alleles existing in an accession (2 × homozygous mutation burden + heterozygous mutation burden)^15,39^.

### Comparison of observed and expected mutation burden under HWE

A bootstrap approach (with replacement) was used to resample cultivated cassava accessions from both Latin American (24 samples) and African (174 samples) breeding pools. The process was repeated for 10,000 times to generate the distribution of expected homozygous and heterozygous mutation burden. For each resampling,

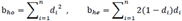

where b_*ho*_ is the expected homozygous mutation burden under HWE, b_*he*_ is the expected heterozygous mutation burden under HWE, n is the total number of deleterious mutations identified (n = 22,495), *d_i_* is the allele frequency of *i*th deleterious allele in the sampled population. The observed mutation burden was calculated for each accession as described in ‘mutation burden in cassava accessions’. The means of observed homozygous and heterozygous mutation were used for the comparison.

#### Data access

Whole genome sequences, raw and imputed HapMapII SNPs can be accessed from CassavaBase at ftp://ftp.cassavabase.org/HapMapII/.

## ACKNOWLEDGEMENTS

This work was supported by the Bill & Melinda Gates Foundation (BMGF: #01511000147), with additional support from NSF Plant Genome Research Project (#1238014) and the UDSA-ARS. We thank Next Generation cassava project (www.nextgencassava.org) for helping us to choose the accessions to include in whole genome sequencing efforts. We thank Simon E. Prochnik (DOE Joint Genome Institute, Walnut Creek, CA, USA) for his timely help during the analysis.

## AUTHORS CONTRIBUTIONS

The manuscript was prepared by P.R., F.L. Data analysis was carried out by P.R., F.L. and E.S.B. Whole genome sequences for 54 accessions included in HapMapl^10^ are provided by J.V.B. W.E., I.Y.R., C.E., R.K. and R.S.B. provided the germplasm for WGS. All authors provided their comments and edited the manuscript. F.L. and E.S.B designed and coordinated the project.

## References

1. Raven, P., Fauquet, C., Swaminathan, M.S., Borlaug, N. & Samper, C. Where Next for Genome Sequencing? Science 311, 468–468 (2006).

2. Olsen, K.M. & Schaal, B.A. Evidence on the origin of cassava: Phylogeography of Manihot esculenta. Proceedings of the National Academy of Sciences 96, 5586-5591 (1999).

3. Wang, W. et al. Cassava genome from a wild ancestor to cultivated varieties. Nat Commun 5(2014).

4. McDonald, M.J., Rice, D.P. & Desai, M.M. Sex speeds adaptation by altering the dynamics of molecular evolution. Nature 531, 233–236 (2016).

5. McKey, D., Elias, M., Pujol, B. & Duputié, A. The evolutionary ecology of clonally propagated domesticated plants. New Phytologist 186, 318–332 (2010).

6. Charlesworth, D. & Willis, J.H. The genetics of inbreeding depression. Nat Rev Genet 10, 783–796 (2009).

7. Rojas, M.C. et al. Analysis of Inbreeding Depression in Eight S1 Cassava Families. Crop Science 49, 543–548 (2009).

8. Nuwamanya, E., Herselman, L. & Ferguson, M. Segregation of selected agronomic traits in six S1 cassava families. Journal of Plant Breeding and Crop Science 3, 154–160 (2011).

9. Allem, A.C. The closest wild relatives of cassava (Manihot esculenta Crantz). Euphytica 107, 123–133 (1999).

10. Bredeson, J.V. et al. Sequencing wild and cultivated cassava and related species reveals extensive interspecific hybridization and genetic diversity. Nat Biotech 34, 562–570 (2016).

11. Prochnik, S. et al. The Cassava Genome: Current Progress, Future Directions. Tropical Plant Biology 5, 88–94 (2012).

12. Browning, Brian L. & Browning, Sharon R. Genotype Imputation with Millions of Reference Samples. The American Journal of Human Genetics 98, 116–126.

13. Chia, J.-M. et al. Maize HapMap2 identifies extant variation from a genome in flux. Nat Genet 44, 803–807 (2012).

14. Tennessen, J.A. et al. Evolution and Functional Impact of Rare Coding Variation from Deep Sequencing of Human Exomes. Science 337, 64–69 (2012).

15. Fu, W. et al. Analysis of 6,515 exomes reveals the recent origin of most human protein-coding variants. Nature 493, 216–220 (2013).

16. Davydov, E.V. et al. Identifying a High Fraction of the Human Genome to be under Selective Constraint Using GERP++. PLoS Comput Biol 6, e1001025 (2010).

17. Rodgers-Melnick, E. et al. Recombination in diverse maize is stable, predictable, and associated with genetic load. Proceedings of the National Academy of Sciences 112, 3823–3828 (2015).

18. Mackay, T.F.C. et al. The Drosophila melanogaster Genetic Reference Panel. Nature 482, 173–178 (2012).

19. Kumar, P., Henikoff, S. & Ng, P.C. Predicting the effects of coding non-synonymous variants on protein function using the SIFT algorithm. Nat. Protocols 4, 1073–1081 (2009).

20. Mezmouk, S. & Ross-Ibarra, J. The Pattern and Distribution of Deleterious Mutations in Maize. G3: Genes/Genomes/Genetics 4, 163–171 (2014).

21. Horvath, P. & Barrangou, R. CRISPR/Cas, the Immune System of Bacteria and Archaea. Science 327, 167–170 (2010).

22. Keller, P.J. & Knop, M. Evolution of Mutational Robustness in the Yeast Genome: A Link to Essential Genes and Meiotic Recombination Hotspots. PLoS Genet 5, e1000533 (2009).

23. Hussin, J.G. et al. Recombination affects accumulation of damaging and disease-associated mutations in human populations. Nat Genet 47, 400–404 (2015).

24. Haddrill, P.R., Halligan, D.L., Tomaras, D. & Charlesworth, B. Reduced efficacy of selection in regions of the Drosophila genome that lack crossing over. Genome Biology 8, 1–9 (2007).

25. Jørgensen, K. et al. Cassava Plants with a Depleted Cyanogenic Glucoside Content in Leaves and Tubers. Distribution of Cyanogenic Glucosides, Their Site of Synthesis and Transport, and Blockage of the Biosynthesis by RNA Interference Technology. Plant Physiology 139, 363–374 (2005).

26. Conn, E.E. Cyanogenic Compounds. Annual Review of Plant Physiology 31, 433451 (1980).

27. Chen, H., Patterson, N. & Reich, D. Population differentiation as a test for selective sweeps. Genome Research 20, 393–402 (2010).

28. Ros, R., Mufioz-Bertomeu, J. & Krueger, S. Serine in plants: biosynthesis, metabolism, and functions. Trends in Plant Science 19, 564–569 (2014).

29. Benstein, R.M. et al. Arabidopsis Phosphoglycerate Dehydrogenase! of the Phosphoserine Pathway Is Essential for Development and Required for Ammonium Assimilation and Tryptophan Biosynthesis. The Plant Cell 25, 50115029 (2013).

30. Marsden, C.D. et al. Bottlenecks and selective sweeps during domestication have increased deleterious genetic variation in dogs. Proceedings of the National Academy of Sciences 113, 152–157 (2016).

31. Agrawal, A.F. & Whitlock, M.C. Inferences About the Distribution of Dominance Drawn From Yeast Gene Knockout Data. Genetics 187, 553–566 (2011).

32. Hohenlohe, P.A. et al. Population Genomics of Parallel Adaptation in Threespine Stickleback using Sequenced RAD Tags. PLoS Genet 6, e1000862 (2010).

33. Danecek, P. et al. The variant call format and VCFtools. Bioinformatics 27, 21562158 (2011).

34. Bradbury, P.J. et al. TASSEL: software for association mapping of complex traits in diverse samples. Bioinformatics 23, 2633–2635 (2007).

35. Consortium, I.C.G.M. High-Resolution Linkage Map and Chromosome-Scale Genome Assembly for Cassava (Manihot esculenta Crantz) from 10 Populations. G3: Genes|Genomes|Genetics 5, 133–144 (2015).

36. Beissinger, T.M., Rosa, G.J., Kaeppler, S.M., Gianola, D. & de Leon, N. Defining window-boundaries for genomic analyses using smoothing spline techniques. Genetics Selection Evolution 47, 1–9 (2015).

37. Du, Z., Zhou, X., Ling, Y., Zhang, Z. & Su, Z. agriGO: a GO analysis toolkit for the agricultural community. Nucleic Acids Research 38, W64–W70 (2010).

38. Supek, F., Bošnjak, M., Škunca, N. & Šmuc, T. REVIGO Summarizes and Visualizes Long Lists of Gene Ontology Terms. PLoS ONE 6, e21800 (2011).

39. Henn, B.M. et al. Distance from sub-Saharan Africa predicts mutational load in diverse human genomes. Proceedings of the National Academy of Sciences 113, E440–E449 (2016).

